# Profile of the digestive and absorptive functions in the intestine of fish with different dietary habits and in response to increased dietary carbohydrate or protein

**DOI:** 10.64898/2026.09.07.749816

**Authors:** Andrew W. Moran, Miran A. Al-Rammahi, Robert L. Bristow, Kieran Magee, Soraya P. Shirazi-Beechey

**Affiliations:** Department of Infection Biology and Microbiomes, Institute of Infection, Veterinary and Ecological Sciences, University of Liverpool, Liverpool, UK; Department of Physiology, Biochemistry and Pharmacology, College of Veterinary Medicine, University of Al-Qadisiyah, Al-Diwaniyah, 58002, Iraq

**Keywords:** SGLT1, B^0^AT1, nutrient absorption, intestines, common carp, Nile tilapia, Rainbow trout

## Abstract

Global aquaculture has grown significantly, but to maintain current consumption, supply must double over the next 20 years. Here, we characterised the digestive and absorptive capacities in the intestines of three commercially important, farmed fish species with varying natural diets: Nile tilapia (*Oreochromis niloticus*), common carp (*Cyprinus carpio*), and rainbow trout (*Oncorhynchus mykiss*). Disaccharidase activity (maltase, sucrase) and sodium-dependent D-glucose absorption were highest in Nile tilapia, followed by common carp, and lowest in rainbow trout. We found regional intestinal differences in disaccharidase activity and SGLT1 expression in the three species. Aminopeptidase A (APA) and aminopeptidase N (APN) activities were typically highest in Nile tilapia. Intestinal expression of the neutral amino acid transporter B^0^AT1 was approx. double in tilapia and carp compared to trout.

We also assessed common carp, zebrafish and Rainbow trout’s digestion profile when fed low-carbohydrate/high protein and high-carbohydrate/low-protein diets. Common carp and zebrafish adapted to high-carbohydrate diets by up-regulating both digestive enzymes and SGLT1. Common carp increased SGLT1 protein abundance 2.7-fold in the foregut and 4.6-fold in the hindgut. Rainbow trout increased disaccharidase activity but not SGLT1 expression in response to a high-carbohydrate diet. Both Rainbow trout and common carp down-regulated their APN activity but upregulated their B^0^AT1 expression 2-fold in response to increased dietary protein. These physiological limitations are critical considerations for formulating sustainable plant-based aquafeeds.

## Introduction

The global expansion of aquaculture is placing increasing pressure on aquafeed production and sustainability. This includes not only increasing production volumes but also ensuring feeds are nutritionally adequate, economically viable, and environmentally sustainable. More recently, the industry has shifted from predominantly marine-based ingredients to more diversified compositions, including plant-derived ingredients. (1,2).

Considering the animal’s physiological and metabolic condition is essential for commercial feed farming, particularly when incorporating ever-increasing plant-based ingredients. These factors can be influenced by species, developmental stage, diet, and the cultivation environment (3).

The fish intestine digests and absorbs food, plays a role in osmoregulation, supports immune function, and acts as a physical barrier against pathogens (4). Studying digestive enzymes and metabolic pathways involved in energy metabolism can help influence the selection of aquafeed ingredients. This can in turn improve nutritional efficiency and the production cycle (3). Little is known about the digestive capacity along the length of the fish gut, even more so about the ability of the fish gut to adapt to an increase in dietary carbohydrate.

Dietary starch is the main digestible carbohydrate in plant feed ingredients (5), and is metabolised in the intestines first by α-amylase to a mixture of oligosaccharides and disaccharides, which are subsequently hydrolysed into monosaccharides by the mucosally-bound disaccharidases, maltase and sucrase (5–7). Glucose and galactose are transported across the brush-border membrane by the active transporter, sodium-glucose co-transport-1 (SGLT1), while fructose is transported by the facilitative transporter, GLUT5 (8–11). Protein is first digested by pepsin (stomach-containing fish only) and trypsin into oligopeptides, which are then hydrolysed into amino acids by the mucosally-bound aminopeptidases (12). Di-and tri-peptides are transported across the brush-border membrane by the active transporter, PEPT1 (13), while amino acids are transported by numerous transporters. One such transporter is B^0^AT1, which transports neutral amino acids across the brush-border membrane (14).

In this paper, we set out to characterise the digestive and absorptive functions of three commercially important farmed fish with different dietary habits: Rainbow trout (*Oncorhynchus mykiss*), common carp (*Cyprinus carpio*), and Nile tilapia (*Oreochromis niloticus*). As of 2022, Nile tilapia was the fourth most globally produced fish and common carp the ninth. Together, Nile tilapia, common carp and Rainbow trout accounted for over 10 million tonnes of fish production globally in 2022, with production coming almost entirely through aquaculture (15). Common carp and zebrafish are both omnivorous species whose guts lack a defined stomach. Carp consume a variety of plant material, invertebrates and detritus. Rainbow trout are carnivores and possess a true stomach and pyloric caeca in their foregut. They consume insects, fish eggs and smaller fish. Nile tilapia also possess a true stomach and have a long intestine. They are herbivores with omnivore tendencies, and their diet consists of phytoplankton, zooplankton, insects, fish parts and detritus (16–18).

Our objectives were (1) to characterise the digestive and absorptive functions in Rainbow trout, common carp and Nile tilapia through measurements of disaccharidase and peptidase activity and the expression of SGLT1 and B^0^AT1. (2) Measure/assay the ability of the common carp and Rainbow trout to adapt to increased carbohydrate or protein in their diet, through the measurement of the disaccharidase and protease activities and the expression of B^0^AT1 and SGLT1. These findings will provide insights into fish capacity to digest and absorb critical components of aquafeeds and support the development of rational diet-formulation strategies.

## Material and methods

### Animals and collection of samples

#### Common carp

Common carp (*Cyprinus carpio*) were sourced from Rodbaston Aquaculture Centre, Staffordshire. Rodbaston uses a recirculation system with a capacity of 45,000 litres. Oxygen is pumped in with the inflow. All tanks are on automatic feeders with the diet containing 36% protein, 7% fat and 45% carbohydrate (Carpco excellent, Coppens International, UK). There is limited usage of cleaning products/disinfectants with 0.5–3.0 tonnes of fish in the system at all times. There is a large k1 moving filter bed and a drum filter, but no ozone or UV treatment.

#### Rainbow trout

Rainbow trout (*Oncorhynchus mykiss*) were sourced from Chirk Trout Farm, Oswestry (CH) and maintained at the University of Liverpool aquarium in a 1000-litre tank on a semi-flow-through system. Trout were fed Skretting commercial feed at 2% of body weight. The temperature of the system was maintained within 11-15 °C.

#### Nile tilapia

Nile tilapia (Oreochromis niloticus) were sourced from the University of Sterling, Sterling, UK. The Nile tilapia were maintained for two weeks at 26-29 °C in a recirculating system consisting of ten 50-litre tanks with a total system volume of approx. 750 litres (approximately 30-40% weekly water change). Tilapia were fed at 2% of body weight on EWOS Micro LR 80P (Cargill, Scotland, UK).

Fish were first anaesthetised by immersion in 100mgl^-^¹ Benzocaine and then sacrificed by pithing of the brain according to UK Home Office Schedule 1 regulations. The average weight and length of carp were 405 g (range ± 184 g) and 22.6 cm (range ± 2.9 cm); whilst trout were 243 g (range ± 51 g) and 24.8 cm (range ± 2.0 cm), and Nile tilapia were 330 g (range ± 56 g) and 22.4 cm (range ± 1.6 cm). Immediately post-mortem, we removed the intestine, opened it longitudinally, and rinsed it in chilled 0.9% (w/v) saline. Depending on the species, the intestinal tissue was separated into either two or three sections. Rainbow trout had their pyloric caeca removed, and the remaining intestine immediately distal to the caeca, labelled midgut, was separated from the latter distal intestine (labelled hindgut) at the point of demarcation between the mid-and hindgut. The intestine of the common carp was cut distal to the intestinal swelling portion of the intestine that resembles a widening of the intestine, and separated into two labelled foregut and hindgut. The intestine of the Nile tilapia was dissected distal to the stomach and separated into three sections, foregut, midgut and hindgut. After rinsing in saline, the intestinal sections were either wrapped in aluminium foil and frozen in liquid nitrogen before storing at −80 °C until use or fixed in 10% neutral buffer formalin (NBF).

### Feeding low/high carbohydrate/protein diets

Common carp and rainbow trout were purchased from Rodbaston Aquaculture Centre and Dunsop Bridge Trout Farm, respectively; whilst zebrafish (AB strain) were bred in the University of Liverpool Aquarium. Carp and trout were maintained in two semi-flow-through systems, each consisting of 10, 50-litre tanks and a total system volume of approx. 750 litres, with temperatures of 12 and 21 °C for trout and carp, respectively. Zebrafish were maintained in a recirculation system in 9 L tanks (stocking density 5 fish/litre), with temperature maintained at 28 ± 1°C, pH 7, and weekly water changes of approximately 20%. All fish were maintained on a 12hr light/12 hrs dark cycle, with UV treatment applied to carp and trout systems. All fish were maintained in good water quality (within desired ranges), with flow-through systems used for carp (approx. 1l/min) and trout (approx. 2-3l/min).

The three fish species were fed a species-specific control diet for 1 week, after which time half the fish were moved onto a species-specific low-carbohydrate/high-protein or high-carbohydrate/low-protein diet for 2 weeks. See table 1 and 2 for the compositions and nutrient analysis of the diets, respectively. The diets were composed so that with low CHO diet was high in protein and the high CHO diet was low in protein allowing the effect of both differing carbohydrate and protein content to be assayed simultaneously. The fish were sacrificed by immersion in 100 mgl^-^¹ benzocaine and then sacrificed by pithing of brain and dissected as above. Due to the much smaller size of the zebrafish intestine, 10 whole zebrafish intestines were pooled per sample number. To consider the influence of circadian periodicity on gene expression (19,20), fish were always killed at 10 am.

**Table 1.** Composition of diets used to assess a change in the carbohydrate and protein content of the diet.

| Nutrient<br>(% of the diet) | Zebrafish |  |  | Common carp |  |  | Rainbow trout |  |  |
| --- | --- | --- | --- | --- | --- | --- | --- | --- | --- |
|  | Carbohydrate |  |  | Carbohydrate |  |  | Carbohydrate |  |  |
|  | Low | Standard | High | Low | Standard | High | Low | Standard | High |
| Fish meal (Danish) | - | - | 3 | - | - | - | - | - | - |
| Fish meal (Organic, white timmings) | 73.3 | 41 | - | 87.5 | 46 | 3 | 70 | 51 | 27 |
| Wheat gluten | 15.8 | 20 | 19.3 | - | 9 | 16.2 | 16 | 15 | 5.1 |
| Corn starch | 9.7 | 30.05 | 59.9 | 11.3 | 37.8 | 60.3 | 7.8 | 22.8 | 52 |
| Vitamins | 0.3 | 0.3 | 0.3 | 0.3 | 0.3 | 0.3 | 0.3 | 0.3 | 0.3 |
| Minerals | 0.4 | 0.4 | 0.4 | 0.4 | 0.4 | 0.4 | 0.4 | 0.4 | 0.4 |
| Binder (CMC) | 0.5 | 0.5 | 0.5 | 0.5 | 0.5 | 0.5 | 0.5 | 0.5 | 0.5 |
| Fish oil (Organic) | - | 7.2 | 16 | - | - | - | - | - | - |
| Fish oil (South American) | - | - | - | - | 3 | 16.2 | - | 5 | 13 |
| Rapeseed oil | - | - | - | - | 3 | - | 5 | 5 | - |
| Lysine | - | 0.55 | 0.6 | - | - | 1.7 | - | - | 1 |
| Leucine | - | - | - | - | - | 0.5 | - | - | 0.4 |
| Arginine | - | - | - | - | - | 0.9 | - | - | 0.3 |
| Total | 100 | 100 | 100 | 100 | 100 | 100 | 100 | 100 | 100 |

**Table 2.** Nutritional analysis of diets used to assess a change in the carbohydrate and protein content of the diet.

| Nutrient | Zebrafish |  |  | Common carp |  |  | Rainbow trout |  |  |
| --- | --- | --- | --- | --- | --- | --- | --- | --- | --- |
|  | Carbohydrate |  |  | Carbohydrate |  |  | Carbohydrate |  |  |
|  | Low | Standard | High | Low | Standard | High | Low | Standard | High |
| Crude protein | 64.71 | 45.70 | 18.70 | 61.48 | 40.07 | 16.08 | 62.55 | 48.49 | 23.58 |
| Crude lipid | 10.76 | 14.21 | 18.21 | 11.50 | 12.94 | 18.23 | 15.33 | 17.90 | 17.31 |
| Carbohydrate | 14.96 | 33.22 | 60.03 | 15.97 | 40.07 | 60.08 | 12.96 | 26.35 | 52.44 |
| Crude ash | 14.51 | 8.34 | 0.98 | 17.15 | 9.24 | 1.03 | 13.88 | 10.23 | 5.57 |
| Crude fiber | 0.09 | 0.11 | 0.11 | 0.01 | 0.06 | 0.09 | 0.09 | 0.09 | 0.03 |
| Arginine | 4.98 | 4.61 | 3.50 | 5.42 | 4.99 | 8.95 | 4.97 | 4.86 | 6.21 |
| Histidine | 1.99 | 1.98 | 1.96 | 1.99 | 1.98 | 1.93 | 1.99 | 1.98 | 1.96 |
| Isoleucine | 3.30 | 3.30 | 3.35 | 3.29 | 3.28 | 3.25 | 3.30 | 3.29 | 3.24 |
| Leucine | 6.07 | 6.15 | 6.46 | 5.95 | 6.03 | 9.35 | 6.08 | 6.09 | 7.64 |
| Lysine | 5.18 | 5.59 | 5.30 | 6.14 | 5.23 | 12.41 | 5.14 | 4.91 | 9.36 |
| Methionine | 0.90 | 1.27 | 3.10 | 1.01 | 1.55 | 3.86 | 1.15 | 1.48 | 3.05 |
| Cystine | 1.00 | 1.19 | 1.81 | 0.74 | 0.97 | 1.77 | 1.01 | 1.06 | 0.95 |
| Phenylalanine | 1.89 | 2.67 | 6.53 | 1.82 | 2.80 | 6.97 | 1.98 | 2.56 | 5.26 |
| Tyrosine | 1.89 | 2.67 | 6.53 | 1.25 | 1.92 | 4.79 | 1.34 | 1.73 | 3.56 |
| Threonine | 2.10 | 2.98 | 7.27 | 2.07 | 3.17 | 7.90 | 2.17 | 2.80 | 5.77 |
| Thryptophan | 0 | 0 | 0 | 0.37 | 0.57 | 1.43 | 0.32 | 0.41 | 0.85 |
| Valine | 2.47 | 3.50 | 8.56 | 1.82 | 2.80 | 6.97 | 1.98 | 2.56 | 5.26 |
| Gross Energy MJ/kg | 23.12 | 23.13 | 22.60 | 22.00 | 21.79 | 21.83 | 23.96 | 23.75 | 21.45 |

### Cloning of SGLT1, PEPT1 and B^0^AT1 mRNA from the rainbow trout and Nile tilapia intestine

RNA was extracted from Rainbow trout and Nile tilapia intestinal tissues (RNeasy Mini Kit, Qiagen), according to the manufacturer’s instructions. The RNA was quantified by UV spectrophotometry [assuming an optical density reading at 260 nm (OD260) that gives a value of 1 = 40 μg/ml] and integrity was determined by gel electrophoresis. Complementary DNA was prepared using random hexamer primers and Superscript III reverse transcriptase (Life Technologies, Paisley, UK), purified using QIAquick PCR Purification Kit (Qiagen, Crawley, West Sussex, UK), and quantified by UV spectrophotometry (assuming an OD260 value of 1 = 33 μg/ml). Consensus primers designed for zebrafish and Salmon SGLT1, B0AT1, and PEPT1 sequences, purchased from Eurogentec (Seraing, Belgium), are listed in Table 3. Polymerase chain reactions were performed using consensus primers (final concentration of 0.2 µM), velocity DNA polymerase (2.5 unit; Bioline, Essex, UK), and 100 ng of cDNA as template in a 25µl reaction. The cycling conditions were an initial denaturation of 94°C for 2 min, followed by 25 cycles of 94°C for 10 s, and annealing for 10 s at 72°C for 30 s. The PCR products were gel purified on 1% agarose gels, cloned into pGEM-T Easy vector (Promega, Southampton, UK), and custom sequenced (Eurofins-MWG, Ebersberg, Germany). Sequence alignments were performed using commercial software (Vector NTI, Life Technologies).

**Table 3.**
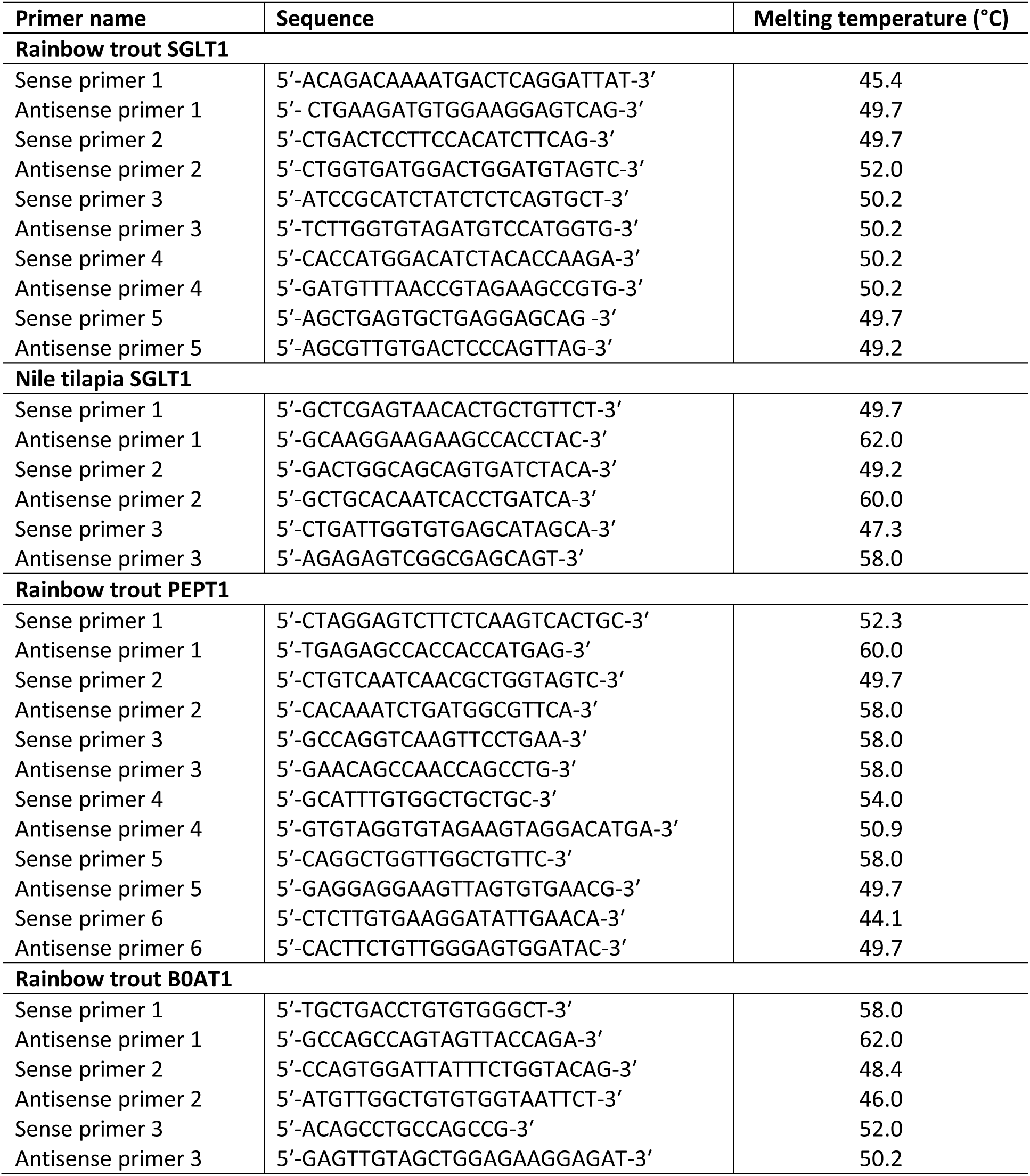
List of primers used in cloning SGLT1, PEPT1 and B^0^AT1 in the rainbow trout and Nile tilapia intestine.

### Preparation of brush-border membrane vesicles

Brush border membrane vesicles (BBMV) were isolated from intestinal tissues based on the procedure described by (21). All steps were carried out at 4°C. Tissues were thawed in a buffer solution (100 mM mannitol, 2 mM HEPES/Tris pH 7.1 with protease inhibitors, 0.5 mM dithiothreitol, 0.2 mM benzamidine, and 0.2 mM phenolmethylsulfonyl fluoride), cut into small pieces and vibrated for 1 min at speed 6 using a FUNDAMIX vibro-mixer (DrM, Dr Mueller AG, Maennedorf, Switzerland) in order to free epithelial cells. Subsequently, the suspension was filtered through a Büchner funnel to remove any muscle and connective tissues. The filtrate was then homogenized using a Polytron (Ystral, Reading, Berkshire, UK) for 30 s at speed 5. Next, MgCl2 was added to the resulting homogenate to a final concentration of 10 mM and the solution stirred on ice for 20 min. The suspension was then centrifuged for 10 min at 3,000 x g (SS34 rotor, Sorvell, UK) and the resulting supernatant was spun for 30 min at 30,000 x g. The pellet was suspended in buffer (100 mM mannitol, 0.1 mM MgSO4, and 20 mM HEPES/Tris pH 7.1) and homogenized with 10 strokes of a Potter Elvehjem Teflon hand-held homogenizer before centrifuging for 30 min at 30,000 x g. The final pellet was re-suspended in an isotonic buffer solution (buffer 3, 300 mM mannitol, 0.1 mM MgSO4, and 20 mM HEPES/Tris pH 7.4) and homogenized by passing through a 27-gauge needle several times. The protein concentration in the BBMV was estimated by its ability to bind Coomassie blue according to the Bio-Rad assay technique (Bio-Rad, Hemel Hempstead, UK). Porcine γ-globulin was used as the standard.

In preparation for western blot analysis, aliquots of freshly prepared BBMV were diluted with sample buffer (62.5 mM Tris/HCl pH 6.8, 10% [v/v] glycerol, 2% [w/v] SDS, 0.05% [v/v] β-mercaptoethanol, 0.05% [w/v] bromophenol blue) and stored at -20°C until use. The remaining BBMV were either diluted 1:100 in buffer 3 and stored at -20°C for enzyme activity determination or were used immediately for glucose uptake studies.

### Enzyme assays

All enzyme assays were carried out at 12, 22 or 28 °C for rainbow trout, common carp and Nile tilapia, respectively.

#### Maltase and Sucrase

Maltase and sucrase activities were measured using the method described previously (21). Diluted BBMV (50 µl) were incubated at species-specific temperatures in 50 µl reaction buffer (56 mM maltose or sucrose and 100 mM Na maleate [pH 6.0]) for 15 min. Reactions were stopped by immersing the tubes in boiling water for 2 min. We measured the concentration of released D-glucose in each reaction mixture using a commercial kit (Roche, Burgess Hill, West Sussex, UK). Specific activity was calculated as nmol/min per mg of brush border membrane protein.

#### Aminopeptidase A and aminopeptidase N

The activity of aminopeptidase A was measured using the α-L-glutamyl-p-nitroanilide (GpNA) as the substrate. A 1.5 mM solution of GpNA was prepared using a buffer consisting of 0.1 M Tris/HCL pH 7, 25 mM CaCl2. A small portion of GpNA was first dissolved into one tenth of the required buffer and dissolved using conc HCl. More reaction buffer was added up to the required amount and brought back to pH 7 using conc NaOH. 50 µl diluted sample was incubated at the appropriate temperature and the reaction started with the addition of 950 µl of assay buffer containing GpNA. The samples were incubated for 30 mins before reading the absorbance against a buffer blank at 410 nm.

The activity of aminopeptidase N was determined using L-leucyl-p-nitroanilide (LpNA) as the substrate. A 100 mM stock solution of LpNA was prepared in EtOH and stored in the dark at 4°C. 50 µL of diluted BBMV was mixed with 935 µl of buffer (51 mM NaH2PO4 pH 7) and incubated at the required temperature. The reaction was started with the addition of 15 µl of substrate and the sample was immediately vibromixed and incubated for 20 mins before reading the absorbance at 410 nm.

### Transport of D-glucose

Na^+^-dependent glucose uptake into BBMV was measured as described (21). The uptake of D-glucose was initiated by the addition of 100 µL of incubation medium (100 mM NaSCN [or KSCN], 100 mM mannitol, 20 mM HEPES/Tris [pH 7.4], 0.1 mM MgSO4, 0.02% [w/v] NaN3 and 0.1 mM [U-^14^C]-D-glucose [10.6 GBq/mmol, Perkin Elmer, Seer Green, Bucks, UK]) to BBMV (100 µg protein) at either 12, 22 or 28°C for rainbow trout, common carp and Nile tilapia, respectively. The reaction was stopped after 3 s by the addition of 1 ml of ice-cold stop buffer (150 mM KCl 20 mM HEPES/Tris [pH 7.4], 0.1 mM MgSO4, 0.02% [w/v] NaN3 and 0.1 mM phlorizin). Aliquots (0.9 ml) of the reaction mixture were removed and filtered under vacuum through a 0.22 µm pore cellulose acetate/nitrate filter (GSTF02500, Millipore, Hertfordshire, UK). We washed the filter with 5 × 1 mL of ice-cold stop buffer, placed it in a vial containing 4 mL of scintillation fluid (Scintisafe 3, Fisher Scientific, UK), and measured the radioactivity retained on the filter using a Tri-Carb 2910TR Liquid Scintillation Analyser (PerkinElmer, Bucks, UK). All uptakes were measured in duplicate.

### Western blotting

We determined the abundance of SGLT1, B^0^AT1, and β-actin proteins in BBMV isolated from fish intestine by western blotting, as described previously (21). Protein components of BBMV were separated by SDS-polyacrylamide gel electrophoresis on 10% (w/v) polyacrylamide mini gels, containing 0.1% (w/v) SDS, and electrotransferred to PVDF membrane (Immun-Blot, Bio-Rad Laboratories Ltd. Hemel Hempstead, UK). Because SGLT1 and B^0^AT1 are similar in size, we prepared separate membranes for each. When blotting for SGLT1 the membranes were blocked by incubating for 1 h at room temperature in PBS-TM buffer (PBS containing 3% (w/v) non-fat dried milk, and 0.3% [v/v] Tween-20) before incubation with SGLT1 antibodies diluted 1:5000 in PBS-TM. The SGLT1 antibody was raised in rabbits (custom synthesis) against a recombinant protein corresponding to amino acids 554-640 of rabbit SGLT1, sharing 54.0% and 51.7% homology with common carp SGLT1 and zebrafish sequences. Immunoreactive bands were detected by incubation for 1 h with affinity-purified horseradish peroxidase-linked anti-rabbit secondary antibody (DAKO Ltd, Cambridge, UK) diluted 1:2000 in PBS-TM, and visualised using Immobilon Western Chemiluminescent HRP Substrate (Millipore, Hertfordshire, UK) and Bio-Max Light Chemiluminescence Film (Sigma-Aldrich, Poole, Dorset, UK).

Blotting for B^0^AT1 was performed using an affinity-purified polyclonal antibody against mouse B^0^AT1 that shares 62.5%, 64.7% and 70.6% homology with rainbow trout, common carp and zebrafish sequences, respectively (NH2-MVRLVLPNPGLEERIC-CONH2, a kind gift from Prof. François Verrey, Universität Zürich, Zürich, Switzerland). A dilution of 1:5000 was used. The PVDF membranes were blocked by incubating them for 3 h at room temperature in TBS-T buffer (TBS containing 0.15% [v/v] Triton X-100) with 5% (w/v) dried skimmed milk powder before incubating them overnight at 4 °C with an affinity-purified B^0^AT1 antibody diluted 1:5000 in TBS-T. We blotted with secondary antibodies and detected immunoreactive bands using the same procedure used for SGLT1.

Before blotting for β-actin, the PVDF membranes were stripped by 3 x 10 min washes in 137 mM NaCl and 20 mM glycine/HCl (pH 2.5), then re-probed with a polyclonal antibody to β-actin (ab8227, diluted 1:5000; Abcam, UK) as a loading control. The antibody was raised to a commercially sensitive peptide from the first 100 residues of the β -actin protein; this region of rabbit β-actin shares 97, 98, 98, and 99% homology to rainbow trout, zebrafish, Nile tilapia and common carp β-actin protein sequences, respectively. The blocking solution consisted of 3% (w/v) skimmed milk powder and 0.3% (v/v) Tween-20 in PBS (PBS-TM).

Incubation and washing buffers were also PBS-TM. Horseradish peroxidase-linked anti-rabbit secondary antibody (DAKO Ltd, Cambridge, UK) diluted 1:2000 in PBS-TM was used and visualised as above.

### Immunohistochemistry

Immunohistochemistry was performed as previously described (22). Tissue sections from rainbow trout, common carp and Nile tilapia small intestine were fixed for 4 hour in 10% neutral buffered formalin (Leica, USA) and then placed in 20% (w/v) sucrose (Fluka, Gillingham, Dorset, UK) in PBS overnight. Subsequently, tissue samples were embedded in OCT embedding medium (Thermoscientific Limited, UK), frozen at -20°C and then kept at -80°C until use. Tissue sections (10µm thick) were sectioned on a cryostat (Leica, CM 1900UV-1-1, Milton Keynes, Buckinghamshire, UK), thaw mounted onto polylysine coated slides and washed five times for 5 minutes each in PBS. Slides were then incubated for 1 hour at room temperature in a humidified chamber with blocking solution [10% (v/v) donkey serum in PBS]. Subsequently, sections were incubated overnight at 4°C with primary polyclonal antibodies. The antibody to SGLT1 was raised (custom synthesis) in rabbits against a synthetic peptide corresponding to amino acids 402–420 (STLFTMDIYTKIRKKASEK; dilution 1:100); the rabbit antibody to mouse B^0^AT1 (see above, dilution 1:200); the antibody to PEPT1 was raised in rabbit to amino acids 140-180 of chicken PEPT1 (dilution 1:100, ab203043, Abcam, Cambridge, UK). After incubation of sections with primary antibodies, slides were washed five times for 5 minutes each in PBS and subsequently stained for 1 hour at room temperature using a 1:500 dilution of Cyanine 3 (Cy3)-or Fluorescein isothiocyanate (FITC)-conjugated anti-rabbit IgG secondary antibodies (Stratech Scientific, Newmarket, UK). The composition of the buffer containing antibodies (primary or secondary) was 2.5% (v/v) donkey serum, 0.25% (w/v) NaN3, and 0.2% (v/v) Triton X-100 in PBS. Finally, slides were washed five times for 5 minutes each in PBS and then mounted with Vectashield Hard Set Mounting Medium with 4ʹ,6-diaminido-2-phenylindole (DAPI) (Vector Laboratories, Peterborough, UK). Immunofluorescent labelling of SGLT1 and B0AT1 proteins was visualised using an epifluorescence microscope (Nikon, Kingston-Upon-Thames, UK), and images were captured with a digital camera (model C4742-96-12G04, Hamamatsu Photonics, Welwyn Garden City, UK). We routinely used omission of primary antibodies as the control.

### Statistical analysis

We tested all parameters for normality using the Shapiro–Wilk test. For comparison of means between different regions of the gut or between different dietary treatments within the same region, a Student’s two-tailed t-test was used to determine statistical significance (GraphPad Prism 11, GraphPad Software Inc., La Jolla, CA). Assessment of the activity and expression along the length of the Nile tilapia gut were performed using an ANOVA and Tukey’s multiple comparison post-test. We set the level of statistical significance at *P* < 0.05.

## Results

### Cloning of SGLT1, B^0^AT1 and PEPT1 in fish intestine

PCR amplicons produced using consensus primers for SGLT1, from RNA extracted from rainbow trout and Nile tilapia intestine, were cloned and sequenced. PCR amplicons produced using consensus primers for B^0^AT1 and PEPT1, from RNA extracted from rainbow trout intestine, were further cloned and sequenced. The resulting sequences were screened against the National Center for Biotechnology Information (NCBI) nonredundant nucleotide database via BlastN. The cloned sequences were homologous to their respective genes in common carp and several mammalian sequences (Fig. 1-3). Rainbow trout SGLT1, PEPT1 and B^0^AT1 sequences were 2280, 2253 and 1934 bp in length, respectively, which translated into peptides of 661, 737 and 635 amino acids, respectively (Fig. 1-3) (Accession No: KY349115 (SGLT1), KY775396 (PEPT1), KY775397 (B^0^AT1). The Nile tilapia SGLT1 sequence was 2013 bp, which translated into a 660-amino-acid peptide (accession No: KY349116). There was high homology in the SGLT1 sequences of the fish species and between the SGLT1 sequences in fish and mammalian species (Fig. 1). Many homologous areas are seen in the B^0^AT1 sequences of fish and mammalian species (Fig. 2), as too with PEPT1; although a 90 bp section of PEPT1 exists where there is almost no homology between rabbit, human, mouse, rainbow trout and common carp (Fig. 3). Importantly, the area of each amino acid sequence used to raise the corresponding primary antibody used in either western blotting or immunohistochemistry show high homology (Fig. 1-3).

**Fig. 1.**
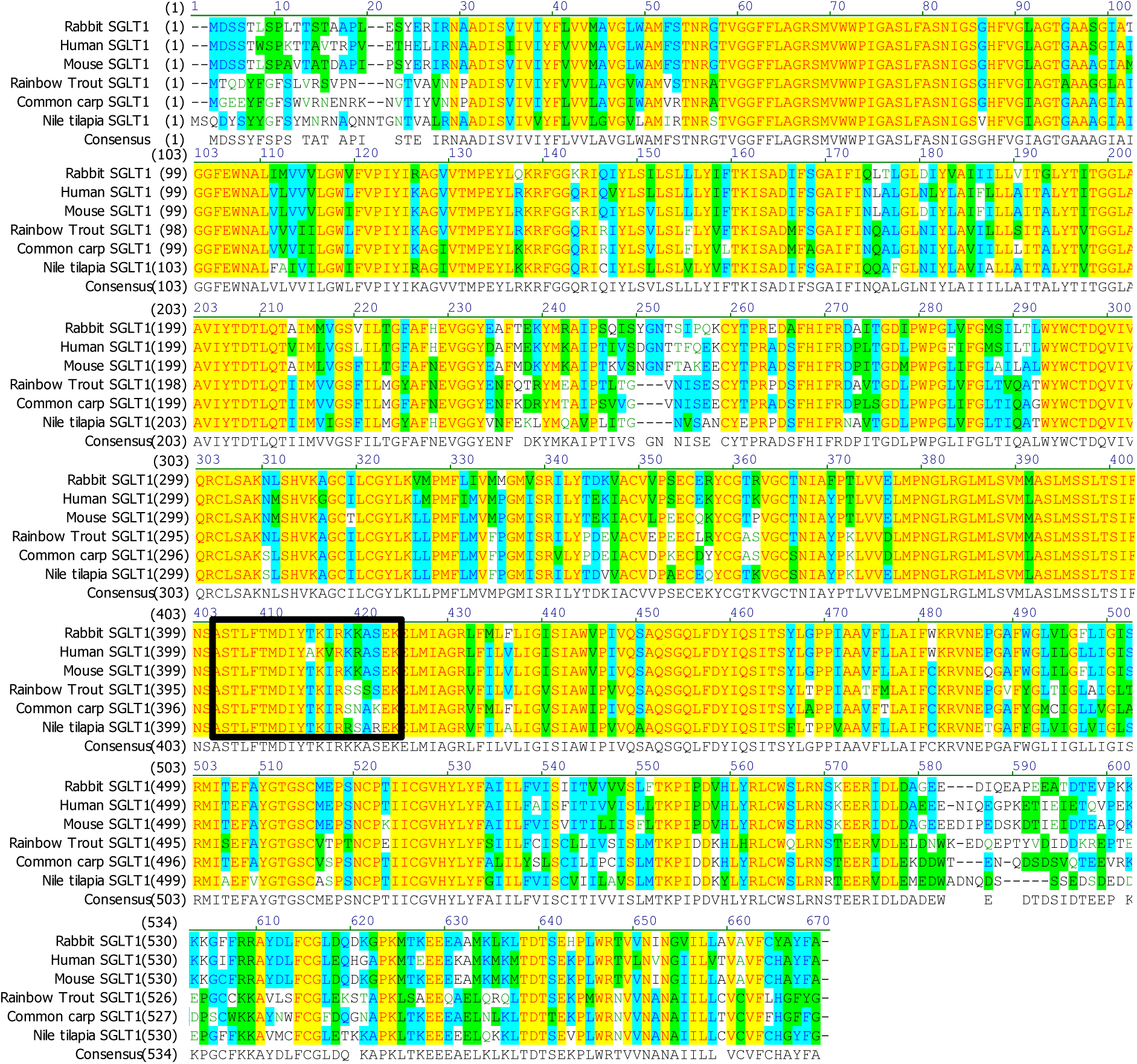
Alignment of rainbow trout and Nile tilapia SGLT1 amino acid sequence (661 and 660 residues, respectively, deduced from cloned mRNA nucleotide sequence) with the corresponding region of rabbit, human, mouse and common carp SGLT1 (numbers in parentheses relate to initiating methionine residues). The black box indicates the rabbit SGLT1 epitope sequence for the SGLT1 antibody used in immunofluorescence.

**Fig. 2.**
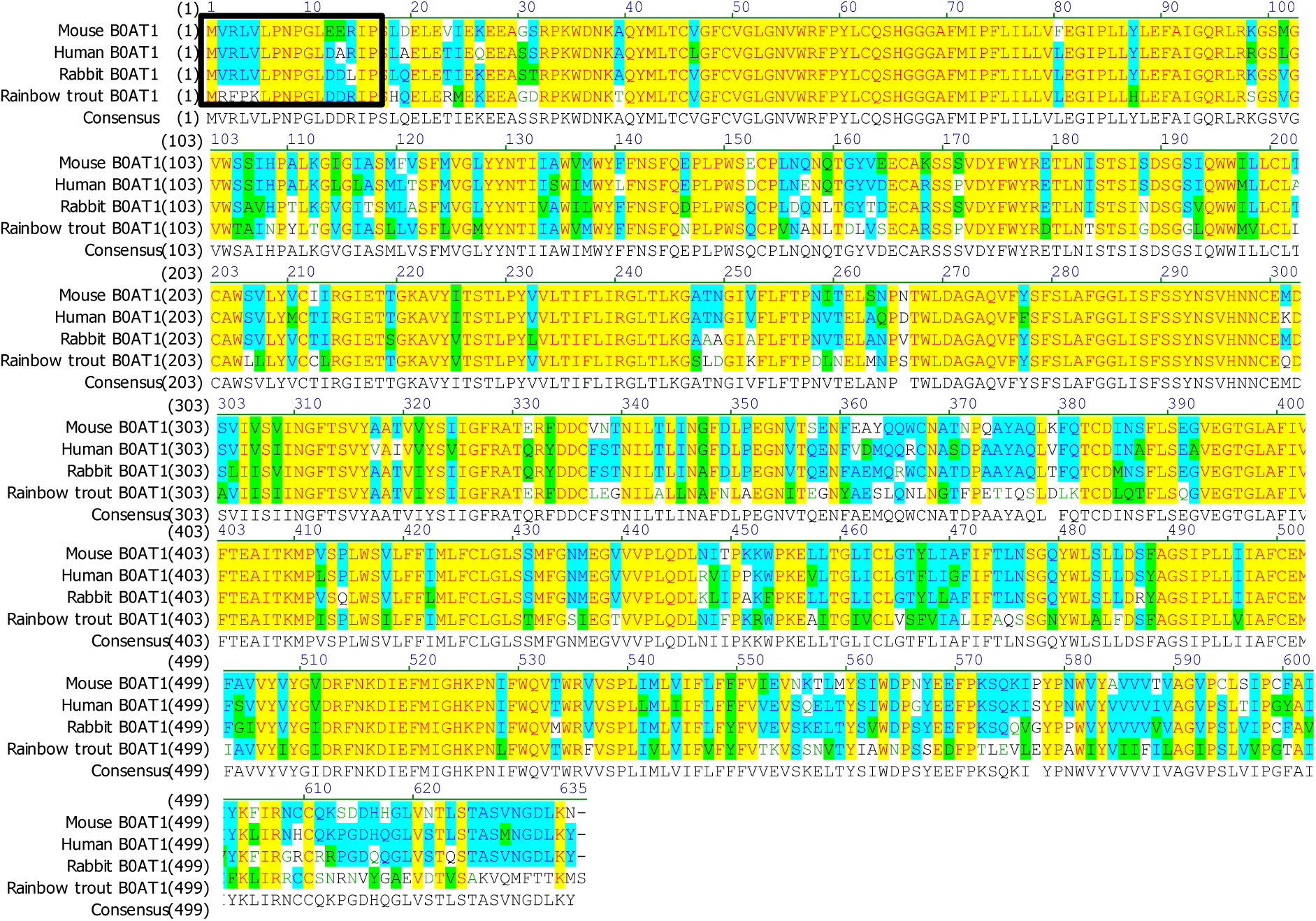
Alignment of rainbow trout B^0^AT1 amino acid sequence (737 residues deduced from cloned mRNA nucleotide sequence) with the corresponding region of mouse, human, and rabbit B^0^AT1 (numbers in parentheses relate to initiating methionine residues). The black box shows the mouse B^0^AT1 epitope sequence for the B^0^AT1 antibody used in western blotting and immunofluorescence.

**Fig. 3.**
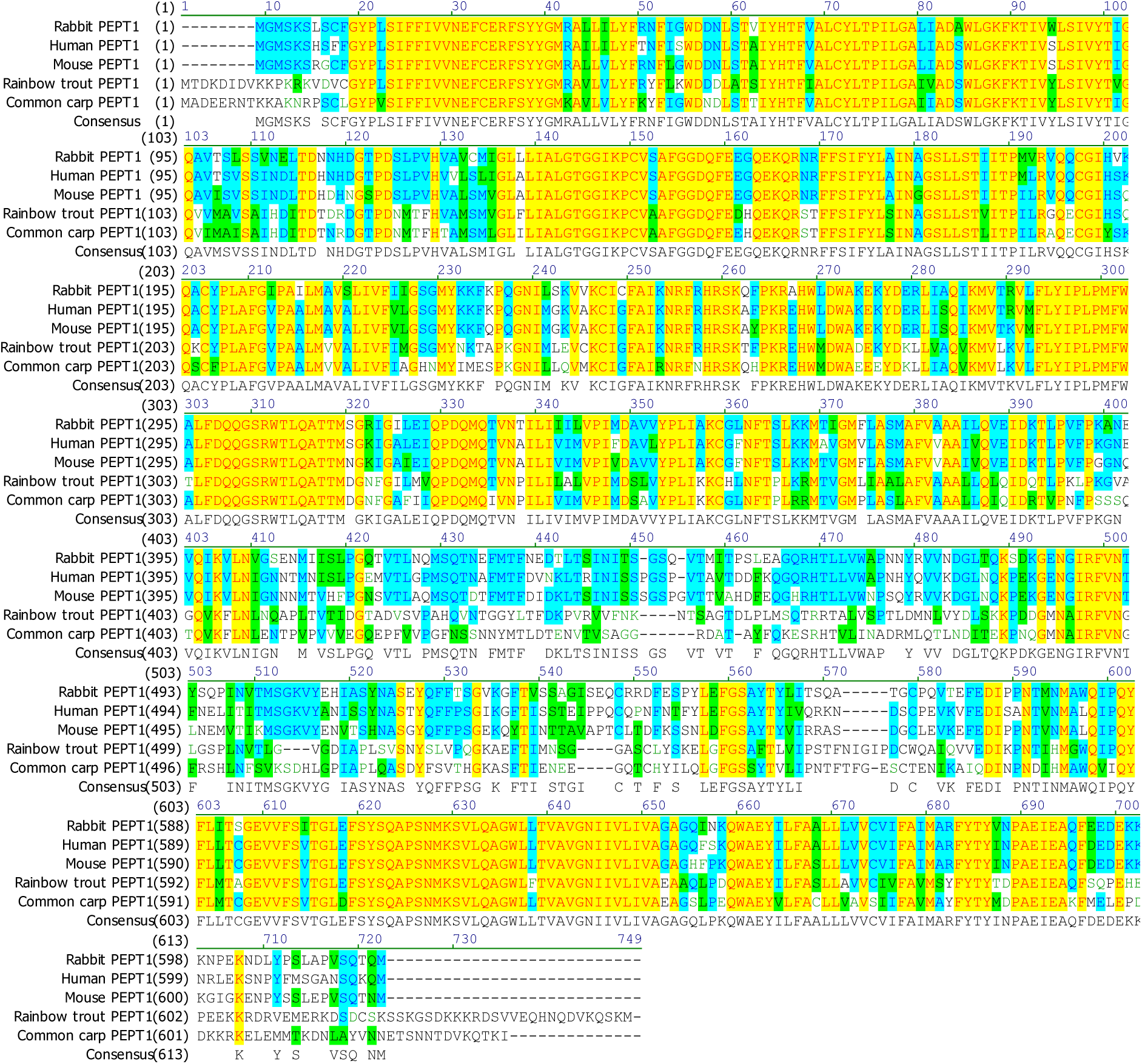
Alignment of rainbow trout PEPT1 amino acid sequence (635 residues deduced from cloned mRNA nucleotide sequence) with the corresponding region of rabbit, human, mouse and common carp PEPT1 (numbers in parentheses relate to initiating methionine residues). The black box indicates the rabbit PEPT1 epitope sequence for the PEPT1 antibody used in immunofluorescence.

### Disaccharidase expression in the intestine of farmed fish

Maltase and sucrase activity were similar in the common carp and Nile tilapia and, on average, were double the activity of maltase and sucrase in rainbow trout (Table 4). Maltase and sucrase activity were similar along the length of the intestine in the common carp. Although rainbow trout showed a similar level of activity in sucrase expression between the midgut and hindgut, there was higher maltase activity in the hindgut of the rainbow trout than the midgut, although this was not significant (74.71 nmon min^-1^ mg^-1^ and 50.56 nmon min^-1^ mg^-1^ in the hindgut and foregut, respectively; *P*=0.1603) (Table 4). Maltase activity in the Nile tilapia gut was approx. double the sucrase activity in all sections of the gut assayed. Sucrase and maltase activity declined along the length of the Nile tilapia gut, with the level of sucrase and maltase activity in the hindgut being 2.6-(P=0.0420) and 11.89-fold (P<0.0001) lower in the hindgut than the foregut, respectively (Table 4).

**Table 4.** Disaccharidase and aminopeptidase activity in the intestine of farmed fish (mean ± SEM)^1^.

| Assay | Activity<br>nmol min<sup>-1</sup> mg<sup>-1</sup> |  |  |  |  |  |  |  |  |  |  |  |  |  |
| --- | --- | --- | --- | --- | --- | --- | --- | --- | --- | --- | --- | --- | --- | --- |
|  | Rainbow trout |  |  |  | Common carp |  |  |  | Nile tilapia |  |  |  |  |  |
|  | Mg |  | Hg |  | Fg |  | Hg |  | Fg |  | Mg |  | Hg |  |
|  | Mean | SEM | Mean | SEM | Mean | SEM | Mean | SEM | Mean | SEM | Mean | SEM | Mean | SEM |
| Maltase | 50.56<sup>a</sup> | 2.45 | 74.71<sup>a</sup> | 10.38 | 102.81<sup>a</sup> | 9.43 | 93.77<sup>a</sup> | 7.18 | 97.26<sup>a</sup> | 13 | 65.32<sup>b</sup> | 5.58 | 8.39<sup>c</sup> | 2.99 |
| Sucrase | 28.28<sup>a</sup> | 2.50 | 26.60<sup>a</sup> | 6.04 | 44.52<sup>a</sup> | 5.57 | 43.65<sup>a</sup> | 3.4 | 50.23<sup>a</sup> | 11.27 | 37.62<sup>ab</sup> | 2.45 | 19.24<sup>b</sup> | 5.94 |
| APA | 52.04<sup>a</sup> | 5.59 | 31.70<sup>b</sup> | 3.60 | 57.63<sup>a</sup> | 6.72 | 13.77<sup>b</sup> | 1.73 | 89.75<sup>ab</sup> | 7.06 | 117.88<sup>a</sup> | 13.20 | 65.19<sup>b</sup> | 11.74 |
| APN | 60.17<sup>a</sup> | 17.91 | 43.93<sup>a</sup> | 13.66 | 163.40<sup>a</sup> | 20.89 | 43.82<sup>b</sup> | 5.37 | 89.53<sup>a</sup> | 12.05 | 97.01<sup>a</sup> | 10.36 | 70.98<sup>a</sup> | 12.62 |

<sup>1</sup>Values are presented as means of six fish. Means in the same row within the same column with different superscripts are significantly different (P < 0.05).

### Aminopeptidase activity in the intestine of farmed fish

Nile tilapia had the highest expression of APA in the intestine, with rainbow trout and common carp displaying smaller and similar levels of expression (Table 4). The region with the highest expression in the Nile tilapia was the midgut, followed by the foregut and then hindgut (1.8-fold decrease between mid and hindgut, *P*=0.0106). APA activity declines along the length of the intestine of both the rainbow trout and common carp, with a 1.6-fold (P=0.0120) and 4.2-fold (P<0.0001) decline in APA activity measured in the hindgut compared to the foregut and midgut of the common carp and rainbow trout, respectively (Table 4).

APN activity was highest in common carp, followed by Nile tilapia and then rainbow trout. Common carp showed a 3.7-fold (*P*=0.0002) decline in APN activity between the foregut and hindgut, whilst similar levels of activity were observed in the rainbow trout along its intestine (Table 4). The activity of APN along the intestine of the Nile tilapia is similar to that of APA, with the highest expression in the midgut followed by the foregut and then hindgut, although there are no significant differences observed (Table 4).

### Transport of D-glucose in the intestine of farmed fish

Overall similar levels of D-glucose transport were measured in the intestine of the common carp and Nile tilapia, conversely a markedly reduction in D-glucose transport in the intestine of the rainbow trout was seen (Fig. 4). The highest rates of D-glucose transport in the rainbow trout were observed in the midgut and a 5.54-fold (*P*< 0.0001) reduction in the rate of D-glucose transport into BBMV from the hindgut was observed compared to the rate of D-glucose transport in the midgut.

**Fig. 4.**
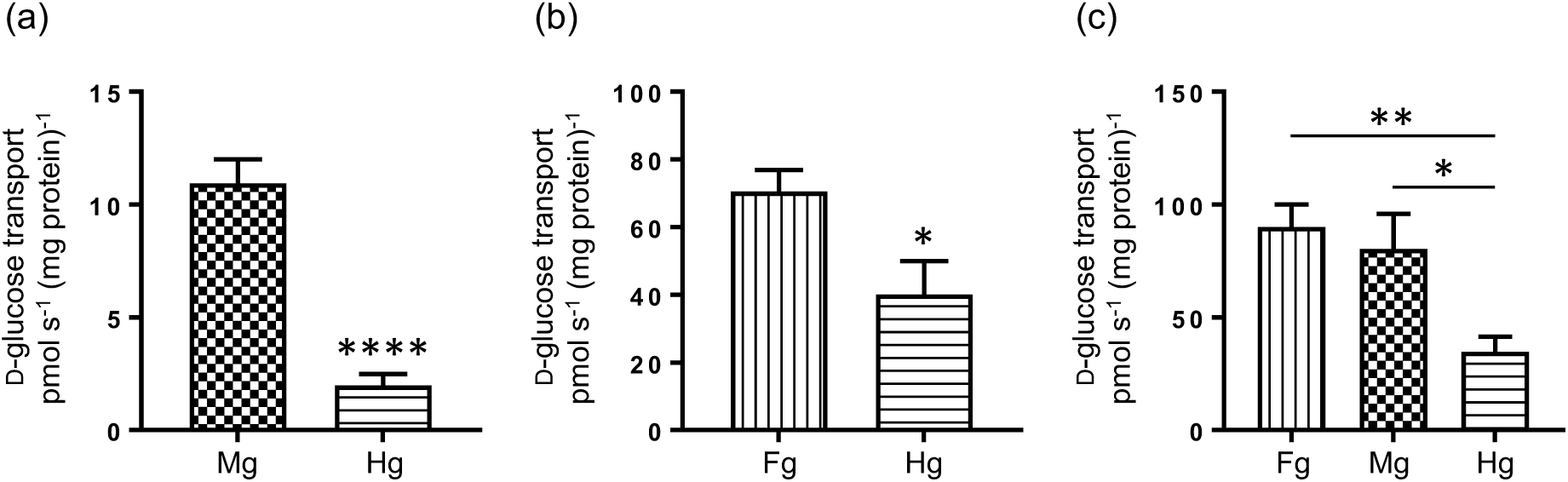
The transport of D-glucose in the intestine of farmed fish. The initial rates of Na^+^-dependent [U^14^C]-D-glucose uptake into BBMV isolated from the foregut (Fg), midgut (Mg) or hindgut (Hg) of rainbow trout (a), common carp (b) or Nile tilapia (c). Data were generated in triplicate. Results are shown as mean ± SEM; n = 7 animals. Statistically significant results were determined using a Student’s two-tailed t-test, where * = P<0.05; ** = P<0.01; and ***= *P*<0.001.

Common carp also displayed higher rates of D-glucose transport in the foregut with a 1.76-fold (*P*=0.0271) reduction in the transport of D-glucose into BBMV isolated from the hindgut compared to BBMV isolated from the foregut (Fig. 4). Nile tilapia showed similar rates of D-glucose transport into BBMV isolated from the foregut and midgut and a 2.6-(*P*=0.0092) and 2.3-fold (*P*=0.0308) reduction in rates of D-glucose transport into BBMV isolated from the hindgut compared to the foregut and midgut, respectively (Fig. 4).

### Expression of B^0^AT1 in the intestine of farmed fish

Common carp and Nile tilapia showed similar levels of expression of B^0^AT1 in the foregut, as measured by western blotting, that were 3.8-(*P*=0.043) and 4.3-fold (*P*=0.0073) higher than the level of B^0^AT1 expression in the midgut of the rainbow trout, respectively (Fig. 5). Common carp showed no change in the expression of B^0^AT1 along its gut contrasting with Nile Tilapia and rainbow trout where no B^0^AT1 was detected in the hindgut (Fig. 5).

**Fig. 5.**
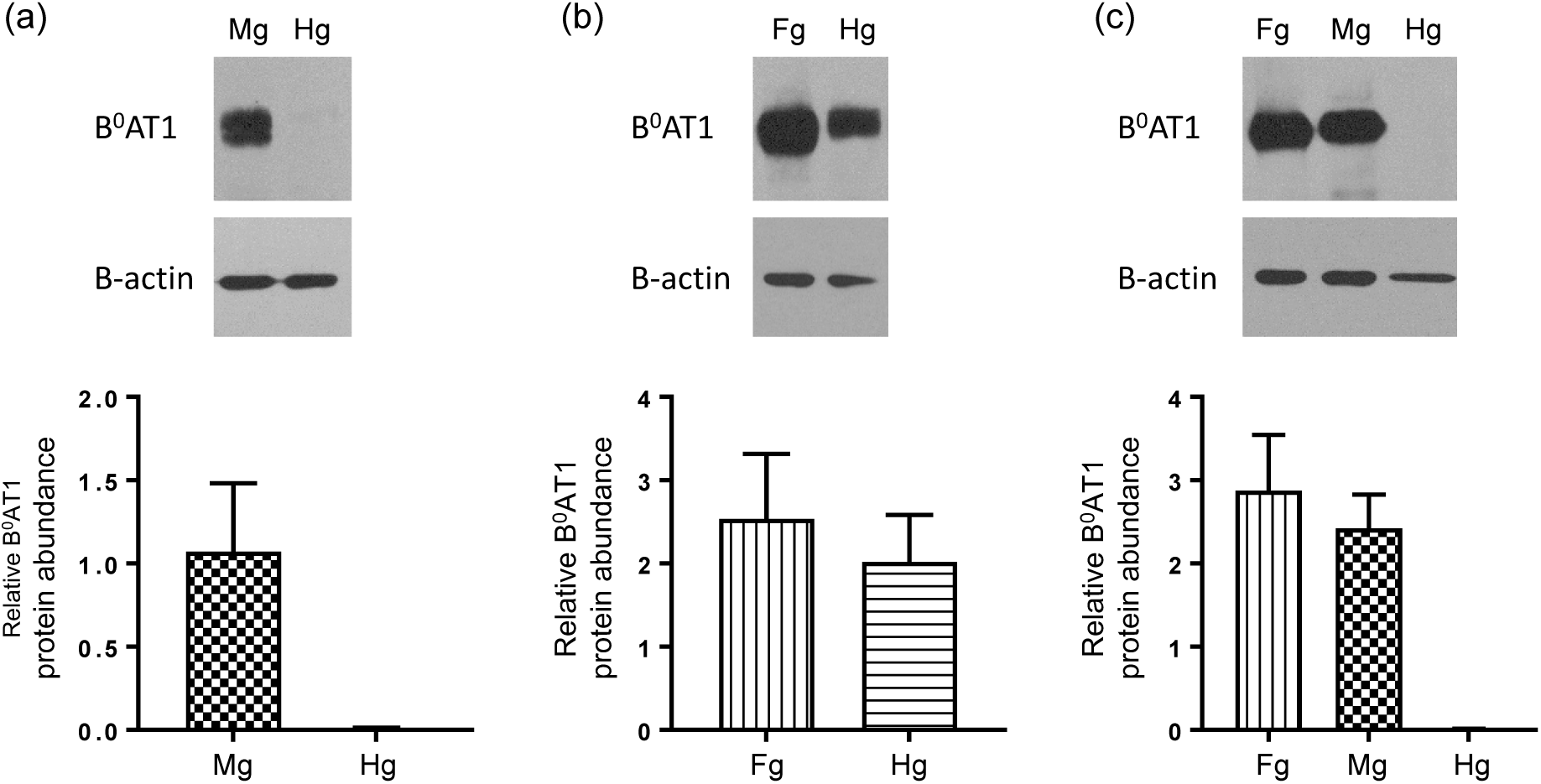
The expression of B^0^AT1 in the intestine of farmed fish. The expression of B^0^AT1 and β-actin proteins in BBMV isolated from the foregut (Fg), midgut (Mg) or hindgut (Hg) of rainbow trout (a), common carp (b) or Nile tilapia (c) were assessed by western blotting (top panel). Densitometric analysis of B^0^AT1 protein abundance normalised to β-actin (bottom panel). Results are shown as mean ± SEM; n = 7 animals.

### Immunohistochemical localisation of SGLT1, B^0^AT1 and PEPT1 in fish intestine

Using antibodies to SGLT1, B^0^AT1 and PEPT1 immunohistochemisty was used to determine the localisation of SGLT1, B^0^AT1 and PEPT1 in the intestine of rainbow trout, common carp and Nile tilapia. In all fish species investigated, SGLT1, B^0^AT1 and PEPT1 showed strong staining on the luminal membrane of all villus cells (Fig. 6), showing a similar pattern of expression to many other species. No basolateral staining on the villus or staining in the underlying layers of the mucosal epithelium was observed.

**Fig. 6.**
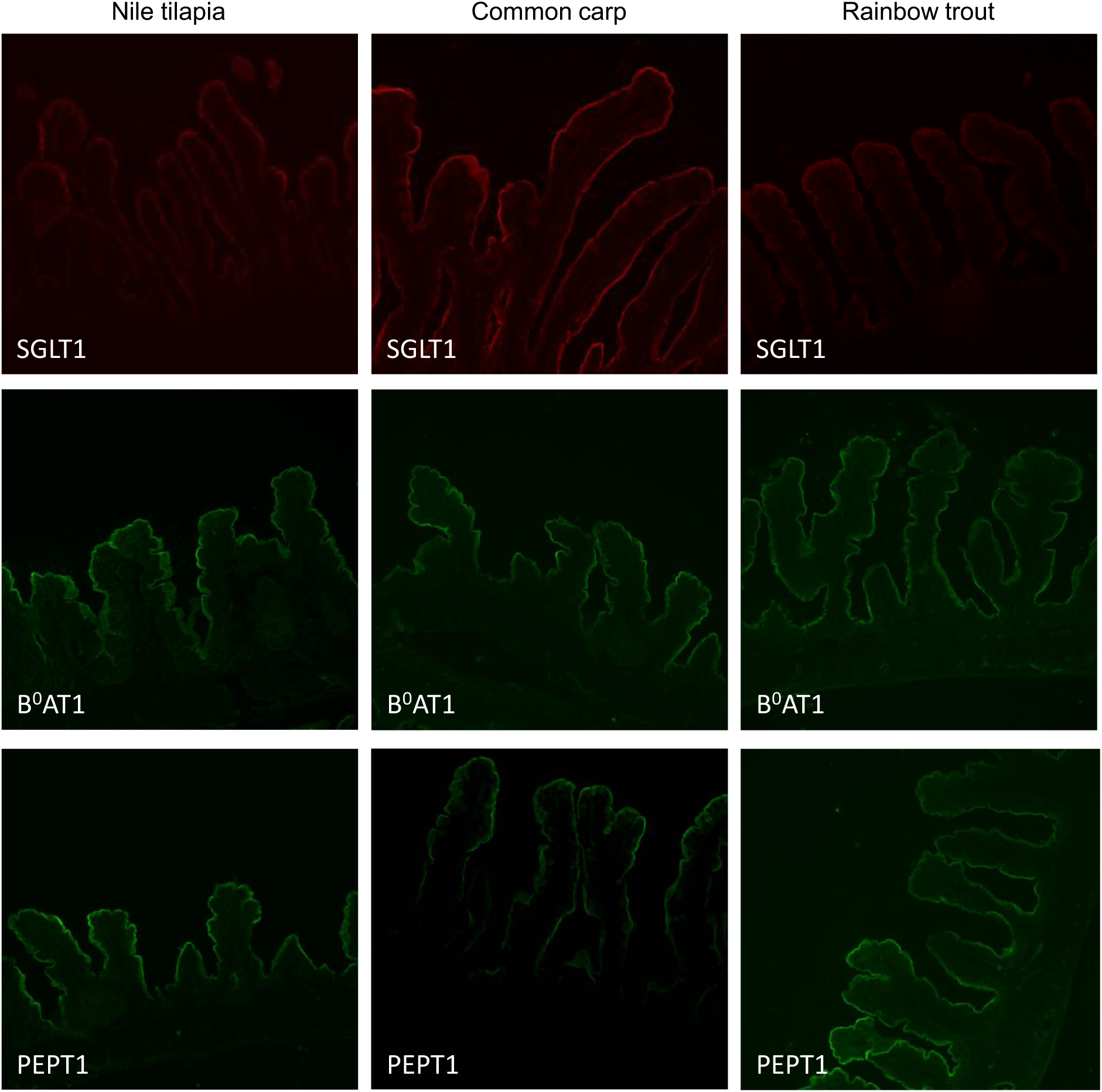
Expression and localisation of SGLT1, B^0^AT1 and PEPT1 in the intestine of Nile tilapia, common carp and rainbow trout. Representative immunofluorescent images show labelling for SGLT1 (red), B^0^AT1 (green) and PEPT1 (green) on the brush border membrane of the entire villus in the intestine of Nile tilapia, common carp and rainbow trout. Sections of rainbow trout midgut and common carp and Nile tilapia foregut were stained with antibodies to SGLT1, B^0^AT1 and PEPT1. Images are 100X magnified.

### Effect of feeding high carbohydrate and high protein diets

Common carp fed the LCHO diet had a mean BW of 69.755 g compared to 49.855 g for common carp fed the HCHO diet (1.4-fold decrease, *P*=0.0345). Similarly, common carp fed the LCHO diet had a body length of 13.29 cm compared to a body length of 11.75 cm for common carp fed the HCHO diet (1.1-fold decrease, *P*=0.0415). There were no differences in either intestinal length or intestinal weight in common carp fed either the LCHO or HCHO diet. There were no differences in either BW or length or intestinal weight or length in rainbow trout fed either the low or high CHO diet. Due to the size and variance of zebrafish, no body or intestinal measurements were taken.

### Disaccharidase and aminopeptidase activity with a change in diet

Rainbow trout that were fed a high CHO diet displayed 1.3-fold (*P*=0.0411) increase in their maltase activity in the midgut compared to rainbow trout fed a low CHO diet (Table 5). No differences in maltase activity were observed in the hindgut of rainbow trout fed either the low or high CHO diet (Table 5). No differences in sucrase activity were observed in any region of the gut of rainbow trout fed either the low or high CHO diet (Table 5). Common carp fed a high CHO diet showed a 2.9-fold (*P*=0.0019) increase in the activity of maltase in their hindgut, compared to carp fed a low CHO diet (Table 5). No differences in maltase activity were observed in the foregut of common carp fed either a low or high CHO diet (Table 5). Sucrase activity in the hindgut of common carp fed a high CHO diet was 2-fold (*P*<0.0001) higher compared to common carp fed a low CHO diet. Sucrase activity in the foregut of common carp fed a high CHO diet was similar to carp fed a low CHO diet (Table 5). Zebrafish fed a high CHO diet displayed a 1.8-fold (*P*<0.0001) increase in maltase activity compared to zebrafish fed a low CHO diet (Table 5); whilst no sucrase activity was detected in either low-or high-CHO-fed zebrafish (Table 5).

**Table 5.** Disaccharidase and aminopeptidase activity in the intestine of fish fed diets of different carbohydrate and protein content (mean ± SEM)^1^.

| Assay | Diet | Activity<br>nmol min<sup>-1</sup> mg<sup>-1</sup> |  |  |  |  |  |  |  |  |  |
| --- | --- | --- | --- | --- | --- | --- | --- | --- | --- | --- | --- |
|  |  | Rainbow trout |  |  |  | Common carp |  |  |  | Zebra fish |  |
|  |  | Mg |  | Hg |  | Fg |  | Hg |  | Mean | SEM |
|  |  | Mean | SEM | Mean | SEM | Mean | SEM | Mean | SEM |  |  |
| Maltase | LC | 8.91 | 1.19 | 4.97 | 0.71 | 15.58 | 2.34 | 2.09 | 0.85 | 19.77 | 1.21 |
|  | HC | 11.98<sup>*</sup> | 0.55 | 5.91 | 0.48 | 20.63 | 2.92 | 16.55<sup>*</sup> | 6.76 | 36.18<sup>*</sup> | 2.05 |
| Sucrase | LC | 2.93 | 0.16 | 2.05 | 0.21 | 3.64 | 0.49 | 3.24 | 0.27 | nd |  |
|  | HC | 2.30 | 0.49 | 2.20 | 0.27 | 4.33 | 0.68 | 6.59<sup>*</sup> | 0.32 | nd |  |
| APA | LP | 29.60 | 2.23 | 41.17 | 1.26 | 48.19 | 7.55 | 41.20 | 6.18 | 20.51 | 1.06 |
|  | HP | 34.75 | 3.99 | 36.49 | 2.61 | 50.17 | 7.61 | 32.95 | 1.67 | 13.67<sup>*</sup> | 0.60 |
| APN | LP | 38.11 | 2.36 | 34.87 | 2.86 | 47.81 | 8.77 | 43.16 | 3.75 | 41.70 | 1.80 |
|  | HP | 40.33 | 2.46 | 27.03<sup>*</sup> | 1.57 | 36.89<sup>*</sup> | 5.73 | 34.25<sup>*</sup> | 1.61 | 21.79<sup>*</sup> | 0.49 |

<sup>1</sup>Values are presented as means of six fish. <sup>\*</sup>Refer to significant differences between either LC and HC or LP and HP within a region of a fish (P < 0.05).

Rainbow trout fed a high-protein diet had similar APA activity in both the midgut and hindgut to rainbow trout fed a low-protein diet (Table 5). Feeding either a low-or high-protein diet to Rainbow trout resulted in similar activity of APN in the midgut. Whereas a 1.3-fold (*P*=0.0375) decrease in APN activity was observed in the hindgut of rainbow trout compared to rainbow trout fed a low-protein diet (Table 5). Common carp fed a high-protein diet had similar APN activity in both the foregut and hindgut to common carp fed a low-protein diet. Conversely, the activity of APN decreased 1.7-fold (*P*=0.0435) and 1.3-fold (*P*=0.0066) in the foregut and hindgut, respectively, in common carp fed a high-protein diet compared to carp fed a low-protein diet (Table 5). Zebrafish fed a high protein diet displayed a 1.5-fold (*P*=0.0002) and 1.9-fold (*P*<0.0001) decrease in APA and APN activity, respectively, when fed a high protein diet compared to zebrafish fed a low protein diet (Table 5).

### SGLT1 and B^0^AT1 expression are altered with changes in diet

Rainbow trout fed a high-protein diet showed no difference in the expression of B^0^AT1 in the midgut compared to trout fed a low-protein diet. However, there was a 2.1-fold (*P*=0.0100) increase in the expression of B^0^AT1 in the hindgut of rainbow trout fed the high protein diet compared to fish fed the low protein diet (Fig. 7). Common carp fed a high protein diet showed a 2-fold (*P*=0.0215) increase in the expression of B^0^AT1 in the foregut compared to the foregut of carp fed the low protein diet (Fig. 7). No differences in the hindgut of common carp fed either the low or high protein diet were seen (Fig. 7). Conversely in zebrafish there was a 2.2-fold (*P*=0.0339) decrease in the expression of B^0^AT1 in the intestine of zebrafish fed a high protein diet compared to fish fed a low protein diet (Fig. 7).

**Fig. 7.**
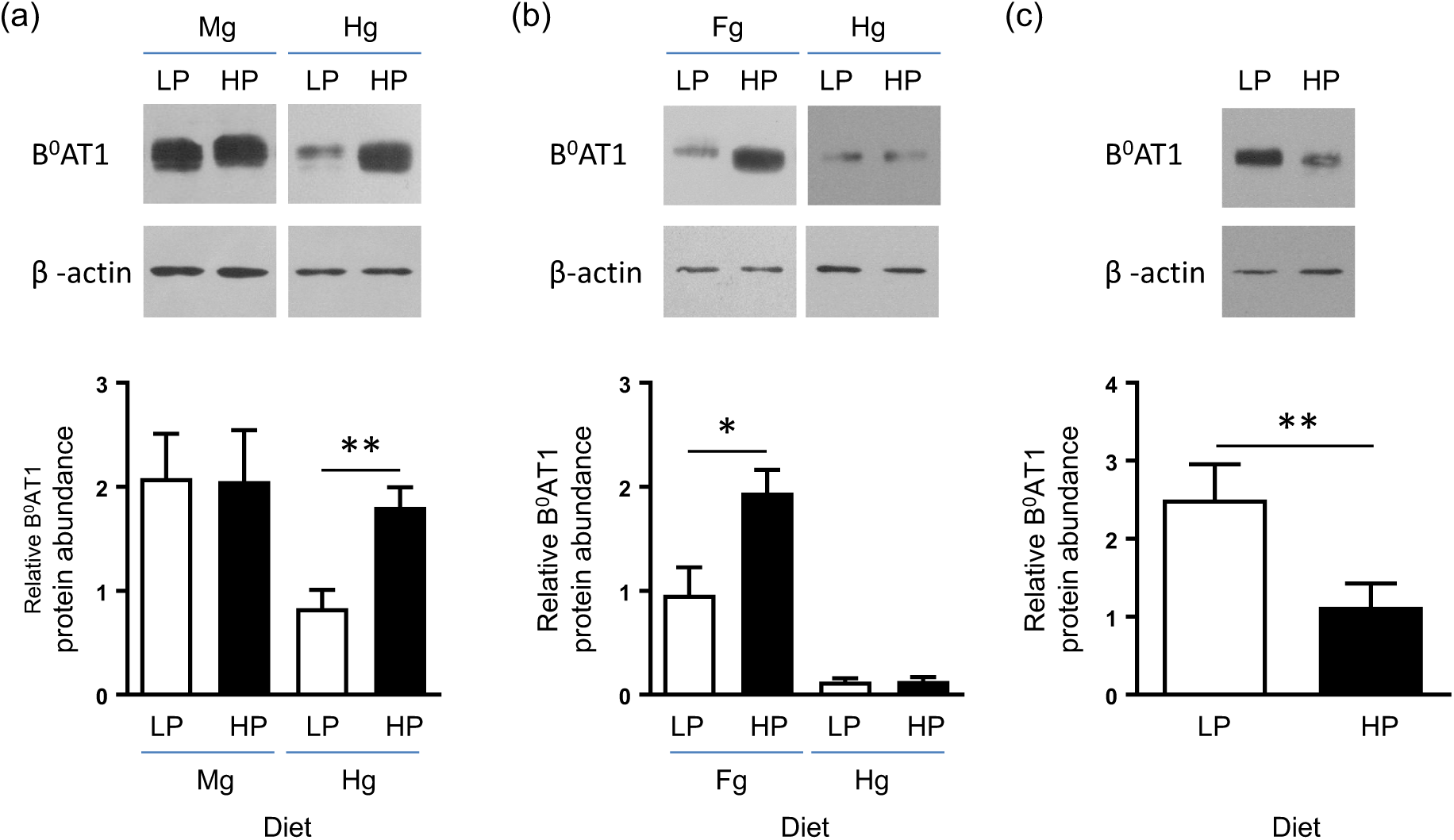
The expression of B^0^AT1 changes with protein content of the diet. The expression of B^0^AT1 and β-actin proteins in BBMV isolated from the foregut (Fg), midgut (Mg) or hindgut (Hg) of rainbow trout (a), common carp (b) or zebrafish (c) fed either a low (LP) or high (HP) protein diet were assessed by western blotting (top panel). Densitometric analysis of B^0^AT1 protein abundance was normalised to β-actin (bottom panel). Results are shown as mean ± SEM; n = 6 animals. Statistically significant results were determined using a Student’s two-tailed t-test, where * = *P*<0.05 and ** = *P*<0.01.

In the intestine of zebrafish fed a high CHO diet, there was a 3.1-fold (*P*=0.003) increase in the abundance of SGLT1 protein, as measured by western blotting, compared to fish fed a low CHO diet. This correlated with a 1.8-fold (*P*=0.0471) increase in the rate of D-glucose transport into BBMV isolated from the intestine of zebrafish fed a high CHO diet compared to fish fed a low CHO diet (Fig. 8). Common carp fed a high CHO diet showed a 2.7-fold (*P*=0.0025) increase in the abundance of SGLT1 protein, as measured by western blotting, in the foregut compared to carp fed a low CHO diet (Fig. 8). There was a larger 4.6-fold (*P*<0.0001) increase in SGLT1 protein in the hindgut of carp fed a high CHO diet compared to carp fed a low CHO diet (Fig. 8). There were no differences in the rates of D-glucose uptake into BBMV isolated from the midgut of rainbow trout fed either the low of high CHO diet. Neither were there any differences in the rates of D-glucose uptake in the hindgut of rainbow trout fed the low and high CHO diet (Fig. 8).

**Fig. 8.**
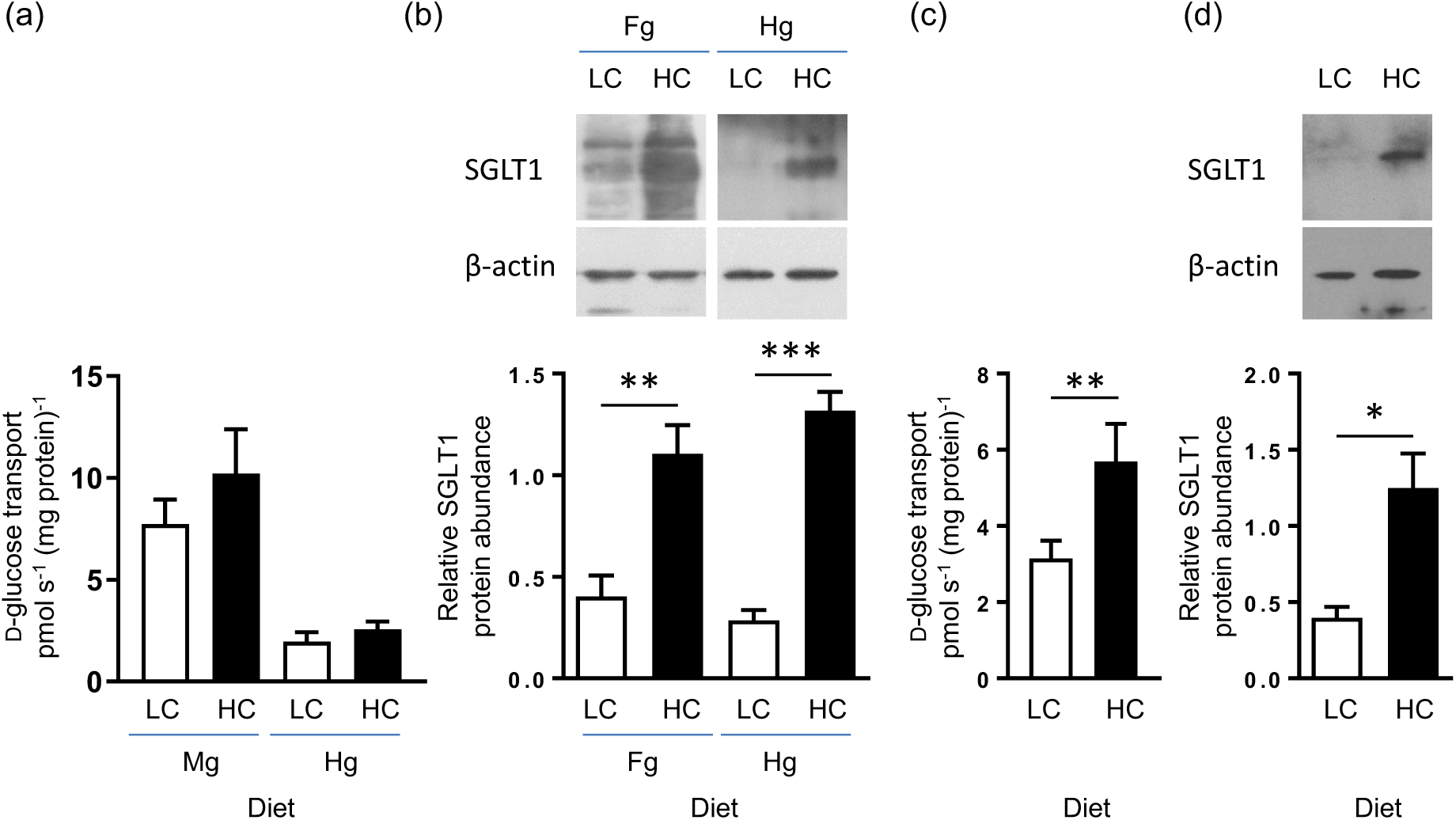
The expression of SGLT1 changed with carbohydrate content of the diet. The expression of SGLT1 was determined in the foregut (Fg), midgut (Mg) or hindgut (Hg) of rainbow trout (a), common carp (b) or zebrafish (c, d) fed either a low (LC) or high (HC) carbohydrate diet. The initial rates of Na^+^-dependent [U^14^C]-D-glucose uptake into BBMV isolated from the midgut (Mg) or hindgut (Hg) of rainbow trout (a) or the whole intestine of zebrafish (c); with data generated in triplicate. SGLT1 and β-actin proteins in BBMV isolated from the foregut (Fg) or hindgut (Hg) of common carp (b) or the whole intestine of zebrafish (d) were assessed by western blotting (top panel). Densitometric analysis of SGLT1 protein abundance was normalised to β-actin (bottom panel). Results are shown as mean ± SEM; n = 6 animals. Statistically significant results were determined using a Student’s two-tailed t-test, where * = P<0.05; ** = P<0.01; and ***= *P*<0.001.

## Discussion

Changes to aquafeed formulations often increase the diet’s carbohydrate content when alternative protein sources are used. Therefore, understanding the effects of increased carbohydrate and the tolerance of the fish intestine is critical, especially in omnivorous and carnivorous fish. Our aim was to better understand carbohydrate and protein digestion and absorption in three fish species with different dietary habits and requirements that account for a significant part of globally commercially farmed fish.

### Assessment of carbohydrate and protein digestion and absorption in the intestine of fish with different dietary habits

There was a clear trend across the three species for carbohydrate digestion and absorption, with Nile tilapia (omnivore/herbivore) having higher rates of maltase and sucrase activity and D-glucose absorption, followed by common carp (omnivore) and then Rainbow trout (carnivore) (Fig. 4, Table 5). As the length of the intestine differs depending on the diet, being shorter than the body length in Rainbow trout and considerably longer than the body length in Nile tilapia, Nile tilapia has a considerable advantage over Rainbow trout in its longer transit time and total carbohydrate digestive activity. The disaccharidase activity and glucose absorption in the common carp and Nile tilapia followed the same intestinal regional expression. Maltase and sucrase activities remained constant along the gut in the common carp and decreased aborally in the Nile Tilapia. The rates of D-glucose absorption decreased aborally in both the common carp and Nile tilapia. The wild Ballan wrasse (Labrus bergylta), which also has a stomachless digestive tract, shows a similar pattern, with maltase expression decreasing aborally (21).

Conversely, maltase activity increased along the intestine in Rainbow trout, sucrase activity remained the same and D-glucose absorption decreased along the gut. Our data on glucose absorption rates in Nile tilapia and Rainbow trout are comparable to those of Subramaniam et al., (2019) who reported higher glucose absorption in Nile tilapia than in rainbow trout, with both species showing intestinal region-specific transport kinetics for glucose absorption (24). Interestingly, Toledo-Solis et al,. (2023) showed that there was a significant decrease in amylase activity in rainbow trout that were fed a vegetable meal replacement diet compared to fish fed a standard fish meal diet (25). Emphasising the need to fully understand the digestive and absorption functions of the fish being tested before replacing seemingly unrelated dietary components.

APA and APN activity and B^0^AT1 expression all showed decreased longitudinal expression in the intestine of the Rainbow trout. This was similar to the pattern of expression in the common carp, where APA and APN activities decreased in the hindgut, and B^0^AT1 expression remained constant along the intestine. This is similar to APN expression in the grass carp intestine, which also decreased aborally. (26). Conversely, APA and APN activities were highest in the mid gut in Nile Tilapia, while B0AT1 expression decreased in the hind gut. Dietary regulation of APN.

Peptidase activity and B^0^AT1 expression were not coordinated with diet, with Nile tilapia having the highest APA and APN activity, followed by the common carp and then Rainbow trout. B^0^AT1 expression was approximately double that in the Nile tilapia and common carp compared to the Rainbow trout. We did not examine disaccharidase and peptidase activity in the pyloric caeca of Rainbow trout, which is a limitation of our study. Atlantic salmon (*Salmo salar*) have a digestive tract similar to Rainbow trout, possess pyloric caeca, and showed 4-fold higher peptidase activity in the pyloric caeca compared to the mid or hind gut (27). We therefore cannot rule out Rainbow trout also having higher disaccharide and peptidase activity in their pyloric caeca, and this is certainly an area for future study.

### Assessment of carbohydrate and protein digestion and absorption with increased carbohydrate and protein in the diet

To assess the effect of increased dietary carbohydrate or protein, fish were brought into the facility and initially maintained on a control diet for 7 days before moving to either a low-carbohydrate/high-protein or high-carbohydrate/low-protein diet for 14 days. The diets were species-specific and isocaloric (Tables 1 & 2)

Disaccharidase activity tended to increase across all species with increased dietary carbohydrate, whereas protease activity was either static or tended to decrease across all species with increased dietary protein. Furthermore, the neutral amino acid transporter, B^0^AT1, expression increased in the intestine of both rainbow trout and common carp fed a high-protein diet. We did not assay PEPT1 levels in the intestine in relation to increased dietary protein, which is responsible for the absorption of di-and tri-peptides and has been shown to be up-regulated by increased dietary protein in rats, rabbits, chickens, Rainbow trout and common carp (28,29).

SGLT1 expression was not increased in the Rainbow trout intestine, but an approx. 2-fold increase was seen in all regions of the common carp and zebrafish intestine when they were fed the high-carbohydrate diet compared to fish fed the low-carbohydrate diet. Increases in activity of disaccharidases and SGTL1 expression occurred along the length of the intestine in the common carp but were restricted to the midgut in Rainbow trout.

At first, it appears that Rainbow trout lack the ability to upregulate carbohydrate digestion and absorption in their gut. However, Polakof & Soengas, (2013) reported a 2-fold increase in SGLT1 mRNA in the midgut of rainbow trout following oral glucose administration. The authors demonstrated a dose-dependent increase in SGLT1 mRNA levels in a rainbow trout *in vitro* model using increasing concentrations of glucose, galactose and mannose (30). In our study, we used complex carbohydrates for the high-carbohydrate diet, whereas Polakof & Soegnas, (2013) used monosaccharides, circumventing the need for amylase, which is expressed at relatively low levels in the Rainbow trout gut compared to omnivorous and herbivorous species (31). Moreover, when Geurden et al., (2007) fed juvenile Rainbow trout a high-carbohydrate diet in the form of dextrin, they saw no upregulation of SGLT1 in the intestine (32). They did, however, report an increase in maltase activity, further strengthening the argument that the low levels of amylase in the gut of Rainbow trout are the limiting factor for carbohydrate-induced increase in SGLT1 expression (32).

Qin et al., (2018) reported increased expression of both SGLT1 and GLUT2 in the intestine of common carp fed a high-carbohydrate meal. They showed a dose-dependent increase in expression, with 40% dietary carbohydrate as the minimum level required to upregulate both SGLT1 and GLUT2 (19). We did not assay GLUT2 levels in this study, but we have previously shown that increased dietary carbohydrate regulates intestinal SGLT1 and GLUT2 expression in pigs, mice, and horses (21,22,33–35).

We show here that both carp and zebrafish are able to upregulate their expression of SGLT1 with increased dietary carbohydrate. In mammals, the upregulation of apically located SGLT1 and basally located GLUT2 in the intestine occurs through the sweet-taste receptor expressed in enteroendocrine cells, located throughout the gut (33,35). We have also assessed the gut’s ability to upregulate SGLT1 expression in response to the inclusion of non-nutritive sweeteners in the diets of mice, pigs, and rabbits (35,36,37). This pathway also involves secretion of the gut hormone GLP-2 (34). Fish have been shown to express the sweet-taste receptor; however, because multiple whole-genome duplication events occurred throughout their evolution, fish contain multiple copies of each STR subunit, with the number differing among species. Yuan et al., (2020) reported a large variance in the number of *Tas1R2* genes in the genomes of fish with European seabass (*Dicentrarchus labrax*), Zebrafish (*Danio rerio*) and common carp (*Cyprinus carpio*) having two copies, Nile tilapia (*Oreochromis niloticus*), medaka (*Oryzias latipes*) and blunt snout bream (*Megalobrama amblycephala*) having three, and grass carp (*Ctenopharyngodon idellus*) having six copies of *Tas1R2* (38).

It is tempting to speculate that fish also regulate SGLT1 expression in the intestine via the STR, allowing dietary manipulation of SGLT1 expression as a strategy for the aquaculture industry. In particular, would Rainbow trout fed non-nutritive sweeteners upregulate the expression of SGLT1, thereby bypassing the low levels of amylase that prevent trout from maximising increased dietary carbohydrate? However, limited work on characterising fish STRs shows that Zebrafish and medaka fish STRs do not respond to sugars (39).

We characterised disaccharidase and peptidase activities and the expression of SGLT1 and B^0^AT1 in Nile tilapia, common carp, and rainbow trout. Furthermore, we showed that common carp and zebrafish can adapt to either a high-carbohydrate or high-protein diet by upregulating digestive and absorptive functions in the intestine. This will help the aquafeed industry take a more science-based approach to designing feed formulations that improve feed efficiency and fish health.

## Acknowledgements

We thank Dr Iain Young for help and assistance with fish nutrition and maintenance.

## Authorship

S.P.S.-B. is responsible for conception and the design of the research, with A.W.M. designing and performing feeding trials, isolation of BBMV, western blotting, glucose transport assays, enzyme assays and carrying out statistical analysis. M.A. performed immunohistochemistry and cloning of SGLT1, PEPT1 and B0AT1. R.B. was responsible for the fish maintenance and assisted with tissue harvesting and processing. K.S. designed and prepared the diets for the feeding trials. S.P.S.-B. and A.W.M. analysed the data, with A.W.M. writing the paper. All authors have read and agreed to the published version of the manuscript.

## References

1. Saromines CJ, Gisbert E, Reinoso S, Martín MLT, Clavijo MP, & Torrecillas S (2026) Valorization of Mushroom By-Products From *Agaricus bisporus* and *Pleurotus ostreatus* as Novel Feed Ingredients for Rainbow Trout (*Oncorhynchus mykiss*) Diets. Aquaculture Nutrition 2026, 6294426.

2. Hua K, Cobcroft JM, Cole A, Condon K, Jerry DR, Mangott A et al. (2019) The Future of Aquatic Protein: Implications for Protein Sources in Aquaculture Diets. One Earth 1, 316– 329.

3. Retcheski MC, Maximowski LV, Escorsin KJS, De Almeida Rosa Kurosaki JK, Romão S, Bitencourt TB et al. (2023) Yarrowia lipolytica biomass—a potential additive to boost metabolic and physiological responses of Nile tilapia. Fish Physiol Biochem 49, 655–670.

4. Wang J, Lei P, Gamil AAA, Lagos L, Yue Y, Schirmer K et al. (2019) Rainbow Trout (Oncorhynchus Mykiss) Intestinal Epithelial Cells as a Model for Studying Gut Immune Function and Effects of Functional Feed Ingredients. Front Immunol 10, 152.

5. Liang Q, Yuan M, Xu L, Lio E, Zhang F, Mou H et al. (2022) Application of enzymes as a feed additive in aquaculture. Mar Life Sci Technol 4, 208–221.

6. Brun A, Mendez-Aranda D, Magallanes ME, Karasov WH, Martínez Del Rio C, Baldwin MW et al. (2020) Duplications and Functional Convergence of Intestinal Carbohydrate-Digesting Enzymes. Molecular Biology and Evolution 37, 1657–1666.

7. Moran AW, Daly K, Al-Rammahi MA, & Shirazi-Beechey SP (2021) Nutrient sensing of gut luminal environment. Proc Nutr Soc 80, 29–36.

8. Shirazi-Beechey SP (1995) Molecular Biology of Intestinal Glucose Transport. Nutr Res Rev 8, 27–41.

9. Ferraris RP, Choe J, & Patel CR (2018) Intestinal Absorption of Fructose. Annu Rev Nutr 38, 41–67.

10. Wright EM, Martín MG, & Turk E (2003) Intestinal absorption in health and disease— sugars. Best Practice & Research Clinical Gastroenterology 17, 943–956.

11. Koepsell H (2020) Glucose transporters in the small intestine in health and disease. Pflugers Arch - Eur J Physiol 472, 1207–1248.

12. Deraison C & Vergnolle N (2026) Proteases in intestinal health and disease. Nat Rev Gastroenterol Hepatol 23, 6–28.

13. Xu G, Zhao W, & Yu Z (2025) Intestinal epithelial transport of bioactive di/tripeptides through PepT1: Molecular mechanism and influencing factors. Food Chemistry 496, 146851.

14. O’Mara M, Oakley A, & Bröer S (2006) Mechanism and Putative Structure of B0-like Neutral Amino Acid Transporters. J Membrane Biol 213, 111–118.

15. (2024) The State of World Fisheries and Aquaculture 2024. FAO.

16. Guillaume, J., Kaushik, S., Bergot, P., and Metailler, R. (2001) Nutrition and Feeding of Fish and Crustaceans. 1st ed., Springer-Verlag, London.

17. Day RD, German DP, Manjakasy JM, Farr I, Hansen MJ, & Tibbetts IR (2011) Enzymatic digestion in stomachless fishes: how a simple gut accommodates both herbivory and carnivory. J Comp Physiol B 181, 603–613.

18. Temesgen M, Getahun A, Lemma B, & Janssens GPJ (2022) Food and Feeding Biology of Nile Tilapia (Oreochromis niloticus) in Lake Langeno, Ethiopia. Sustainability 14, 974.

19. Qin C, Yang L, Zheng W, Yan X, Lu R, Xie D et al. (2018) Effects of dietary glucose and sodium chloride on intestinal glucose absorption of common carp (Cyprinus carpio L.). Biochemical and Biophysical Research Communications 495, 1948–1955.

20. Pascon G, Daniso E, Cardinaletti G, Messina M, Campagnolo F, Zuccaccia D et al. (2024) Postprandial kinetics of digestive function in rainbow trout (Oncorhynchus mykiss): genes expression, enzymatic activity and blood biochemistry as a practical tool for nutritional studies. Comparative Biochemistry and Physiology Part A: Molecular & Integrative Physiology 288, 111559.

21. Moran AW, Al-Rammahi MA, Arora DK, Batchelor DJ, Coulter EA, Ionescu C et al. (2010) Expression of Na^+^ /glucose co-transporter 1 (SGLT1) in the intestine of piglets weaned to different concentrations of dietary carbohydrate. Br J Nutr 104, 647–655.

22. Dyer J, Al-Rammahi M, Waterfall L, Salmon KSH, Geor RJ, Bouré L et al. (2009) Adaptive response of equine intestinal Na+/glucose co-transporter (SGLT1) to an increase in dietary soluble carbohydrate. Pflugers Arch - Eur J Physiol 458, 419–430.

23. Zhou W, Krogdahl Å, Sæle Ø, Chikwati E, Løkka G, & Kortner TM (2021) Digestive and immune functions in the intestine of wild Ballan wrasse (Labrus bergylta). Comparative Biochemistry and Physiology Part A: Molecular & Integrative Physiology 260, 111011.

24. Subramaniam M, Weber LP, & Loewen ME (2019) Intestinal electrogenic sodium-dependent glucose absorption in tilapia and trout reveal species differences in *SLC5A* - associated kinetic segmental segregation. *American Journal of Physiology-Regulatory*, Integrative and Comparative Physiology 316, R222–R234.

25. Toledo-Solís FJ, Larrán AM, Ortiz-Delgado JB, Sarasquete C, Dias J, Morais S et al. (2023) Specific Blood Plasma Circulating miRs Are Associated with the Physiological Impact of Total Fish Meal Replacement with Soybean Meal in Diets for Rainbow Trout (Oncorhynchus mykiss). Biology 12, 937.

26. Tang J, Qu F, Tang X, Zhao Q, Wang Y, Zhou Y et al. (2016) Molecular characterization and dietary regulation of aminopeptidase N (APN) in the grass carp (Ctenopharyngodon idella). Gene 582, 77–84.

27. Martínez-Llorens S, Peruzzi S, Falk-Petersen I-B, Godoy-Olmos S, Ulleberg LO, Tomás-Vidal A et al. (2021) Digestive tract morphology and enzyme activities of juvenile diploid and triploid Atlantic salmon (Salmo salar) fed fishmeal-based diets with or without fish protein hydrolysates. PLoS ONE 16, e0245216.

28. Verri T, Barca A, Pisani P, Piccinni B, Storelli C, & Romano A (2017) Di-and tripeptide transport in vertebrates: the contribution of teleost fish models. J Comp Physiol B 187, 395–462.

29. Spanier B & Rohm F (2018) Proton Coupled Oligopeptide Transporter 1 (PepT1) Function, Regulation, and Influence on the Intestinal Homeostasis. In Comprehensive Physiology, 1st edn, pp. 843–869 [Terjung R, editor]. Wiley.

30. Polakof S & Soengas JL (2013) Evidence of sugar sensitive genes in the gut of a carnivorous fish species. Comparative Biochemistry and Physiology Part B: Biochemistry and Molecular Biology 166, 58–64.

31. Hidalgo MC, Urea E, & Sanz A (1999) Comparative study of digestive enzymes in fish with different nutritional habits. Proteolytic and amylase activities. Aquaculture 170, 267–283.

32. Geurden I, Aramendi M, Zambonino-Infante J, & Panserat S (2007) Early feeding of carnivorous rainbow trout (*Oncorhynchus mykiss*) with a hyperglucidic diet during a short period: effect on dietary glucose utilization in juveniles. American Journal of Physiology-Regulatory, Integrative and Comparative Physiology 292, R2275–R2283.

33. Margolskee RF, Dyer J, Kokrashvili Z, Salmon KSH, Ilegems E, Daly K et al. (2007) T1R3 and gustducin in gut sense sugars to regulate expression of Na^+^ -glucose cotransporter 1. Proc Natl Acad Sci USA 104, 15075–15080.

34. Moran AW, Al-Rammahi MA, Batchelor DJ, Bravo DM, & Shirazi-Beechey SP (2018) Glucagon-Like Peptide-2 and the Enteric Nervous System Are Components of Cell-Cell Communication Pathway Regulating Intestinal Na+/Glucose Co-transport. Front Nutr 5, 101.

35. Moran AW, Alrammahi M, Daly K, Weatherburn D, Ionescu C, Blanchard A et al. (2025) Luminal Sweet Sensing and Enteric Nervous System Participate in Regulation of Intestinal Glucose Transporter, GLUT2. Nutrients 17, 1547.

36. Moran AW, Al-Rammahi MA, Arora DK, Batchelor DJ, Coulter EA, Daly K et al. (2010) Expression of Na^+^ /glucose co-transporter 1 (SGLT1) is enhanced by supplementation of the diet of weaning piglets with artificial sweeteners. Br J Nutr 104, 637–646.

37. Moran AW, Al-Rammahi MA, Daly K, Grand E, Ionescu C, Bravo DM et al. (2020) Consumption ofa Natural High-Intensity SweetenerEnhances Activity and Expression of Rabbit Intestinal Na+/Glucose Cotransporter 1 (SGLT1) and Improves Colibacillosis-InducedEnteric Disorders. Journal of Agricultural and Food Chemistry 68, 441–450.

38. Yuan X-C, Liang X-F, Cai W-J, He S, Guo W-J, & Mai K-S (2020) Expansion of sweet taste receptor genes in grass carp (Ctenopharyngodon idellus) coincided with vegetarian adaptation. BMC Evol Biol 20, 25.

39. Oike H, Nagai T, Furuyama A, Okada S, Aihara Y, Ishimaru Y et al. (2007) Characterization of Ligands for Fish Taste Receptors. J Neurosci 27, 5584–5592.

